# Inferring large networks with matrix factorisation to capture non-linear dependencies among genes using sparse single-cell profiles

**DOI:** 10.64898/2026.03.08.710347

**Authors:** Indra prakash Jha, Ankur Gajendra Meshran, Vinodh Kumar, Kedar Nath Natarajan, Vibhor Kumar

## Abstract

Inference of non-linear dependencies among a large number of features from their scores is an unresolved challenge. Especially when the feature-score matrix is very sparse, like single-cell transcriptome profiles, the problem of estimating dependencies among a large number of features to infer a network becomes an even more daunting task. Here, we propose a method of network inference in reduced dimension (NIRD) to handle sparsity and computational complexity while still inferring non-linear dependencies among genes (features) using large, sparse gene-expression matrices (feature scores). Our method is based on matrix factorisation of gene-expression matrix to facilitate internal imputation as well as network inference using tree ensemble-based non-linear regression. NIRD not only outperformed many other methods across multiple single-cell transcriptomic profiles but also provided consistent inferred networks even in the presence of batch effects. The consistency provided by NIRD helps compare inferred networks to identify genuine genes responsible for changes in regulation due to disease or stress. NIRD can also be used with RNA velocity for better inference of non-linear causality. Application of NIRD with RNA-velocity could improve the prediction of direct targets of transcription factors in human embryonic stem cells, which we validated using ChIP-seq and gene-knockout datasets.

## 1. Introduction

Inferring dependency networks is crucial for understanding the underlying dynamics of complex regulatory processes and causality in biological systems. Inferring networks among regulatory elements in genomics has remained an open problem. For example, inferring a large network of direct gene dependencies from transcriptomic profiles has remained in demand and remains an open problem for understanding the regulation of developmental processes and diseases. As regulation among genes often vary according to cell type and cell-state, it becomes a necessity to use single-cell transcriptome profile to infer gene-network. Single-cell transcriptomic profiles enable the capture of heterogeneity among individual cells belonging to a cell-type and untangling the dynamics of cellular differentiation. With conventional bulk RNA-Seq approaches, the blurring effect from multiple cell types often reduces the strength or statistical power of the evidence, hindering clear insights. Although many methods have been proposed for inferring gene regulatory networks from bulk expression profiles, they have not proved suitable for sparse scRNA-seq profiles [1]. Pratapa et al.[2] evaluated 12 network inference methods using experimental and simulated single-cell expression profiles and concluded that the performances of all the tested methods were less than ideal. Pratapa et al. also concluded that GENIE3 and GRNBoost2 were leading and consistent performers[2]. Gene Network Inference with Ensemble of trees(GENIE3) which also appeared as top performer in the DREAM5 network inference challenge[3] is based on ensemble of decision trees [4]. Another method for gene-network inference, GRNBoost2[5] uses gradient boosting based on the GENIE3 approach and it was adopted by the single-cell clustering tool SCENIC [6]. Recently, deep learning based approaches are being applied for gene-regulatory inference, however they need pre-training using a large amount of datasets [7] and further optimization in terms of required computational resources. Recently after performing a large-scale benchmarking using single cell profiles for network inference methods, Chevalley et al.[8] concluded that GRNBoost2 had high recall on biological evaluation. Chevalley et al. concluded that methods to detect non-linear dependencies among genes have better performance even for single-cell transcriptomes. However, most network inference methods do not scale to large networks consisting of many genes. Even, to infer a large network consisting of more than 5000 genes, GENIE3 and GRNboost2 often become computationally demanding. While GENIE3 and GRNBoost2 are computationally intensive, other methods like ARACNE [9]and RELNET [10] based on mutual information are simplified and fast. It is often not trivial to find a balance between time and accuracy in choosing the required method. Hence, there is still scope for better methods to predict direct non-linear dependencies among genes for large-scale network inference with an optimised computational approach.

In this study, we have proposed an algorithm for inferring gene regulatory networks using large, sparse datasets, such as scRNA-seq expression profiles. Our method also leads to a conceptual advancement that can have wider application for modelling large numbers of linear and non-linear dependencies. Our method is based on matrix factorisation, followed by a non-linear regression to estimate factor importance. We have also tried to compare different matrix factorisation-based algorithms for the robustness of the modelling and the predicted network. Further, we have shown how our method can be generalised to RNA-velocity to infer causality through network inference.

## Results

For our approach of network inference in reduced dimension (NIRD) we first apply matrix factorisation on gene expression matrix. The dimension is reduced such that we represent every sample (or single-cell) in terms of basis vectors. During matrix factorisation we also get loading of each gene on the basis vector. Further, for every gene we apply non-linear regression model based on ensemble of decision trees (RandomForest) to predict its expression across samples (or single-cells) using the basis vector. For every gene we estimate the feature-importance for every basis vector (Material and Methods, Fig. 1). We back-project the feature importances of basis vectors to genes based on their contribution (or correlation) with the basis vector. For every gene the strength of interaction with other genes is considered as their estimated back-projected feature importance while modeling its expression (Material and Methods, Fig. 1). We evaluated NIRD method using several bulk and single cell transcriptome datasets.

**Figure 1.**
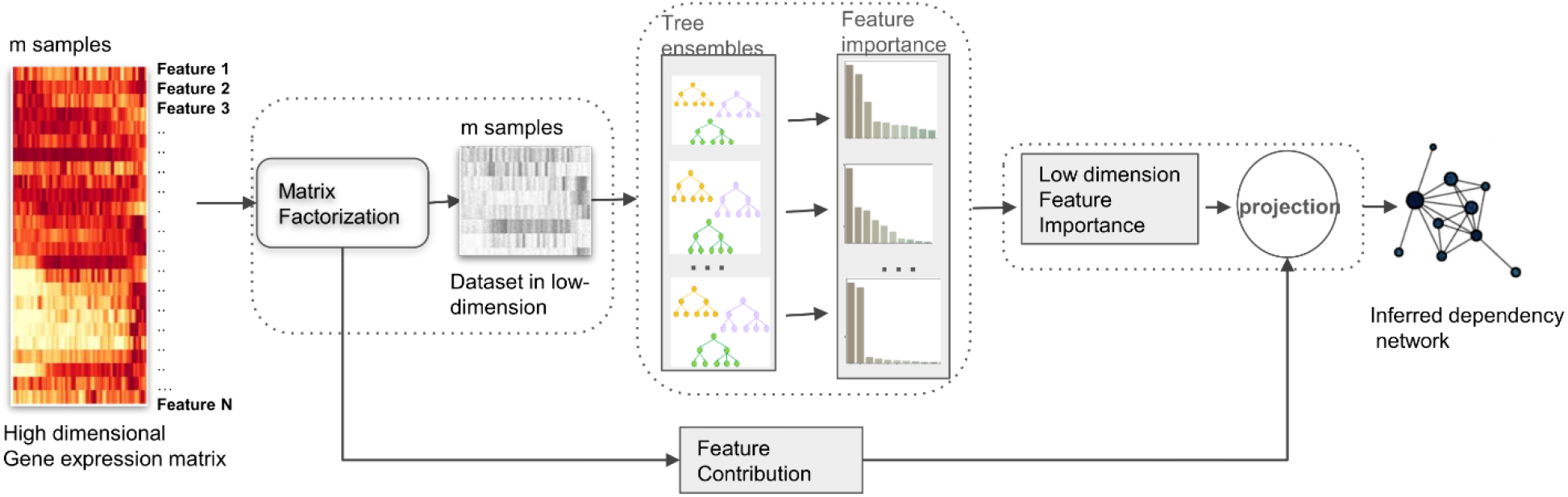
The pictorial view of NIRD approach. In order to find dependency between N features (genes) using their scores (gene-expression), we first perform dimension reduction to achieve a new set of basis vectors and the projections of data-points in them. For each original feature (gene), we model its score (gene expression) using random forests, using projection values onto the basis vectors, and determine the feature importance of different basis vectors in the model. We project the feature importances of each basis vector onto the original genes (features) to estimate their actual dependencies with the target gene (feature). Thus, we obtain a dependency network between features (genes) using our NIRD approach.

### Evaluation and insights using DREAM challenge expression dataset

We used three experimental datasets made available by the DREAM5 challenge consortium as they also have true positive interaction information as Gold set. The three datasets, we used for evaluation are based on the microarray data of gene expression of Escherichia coli (E. coli), Staphylococcus aureus (S. aureus), Saccharomyces cerevisiae (S. cerevisiae). Network inference using original expression profiles revealed that the result using approach of NIRD was comparable or better in comparison to other benchmarked methods GENIE3, GRNboost2, ARACNE and RELNET (Fig. 2A). In order to have statistical insights we used different kinds of matrix factorisation methods and among them NIRD based on principal component analysis (eigen vector decomposition) was consistently better than GRNboost2, GENIE3 and ARACNE (Fig. 2A). Whereas NIRD based on SepNMF matrix factorisation method showed best performance for gene-expression data of S. aureus and E. coli(Fig. 2A). In terms of computation time needed by different methods, GRNboost2 needed the highest amount of computation time. Most NIRD-based methods required less computation time than GENIE3 and GRNboost2 and often outperformed them (Fig. 2B). Overall, using the DREAM5 datasets revealed that NIRD can be a reliable option for good performance at low computation time, even for bulk gene-expression profiles.

**Figure 2.**
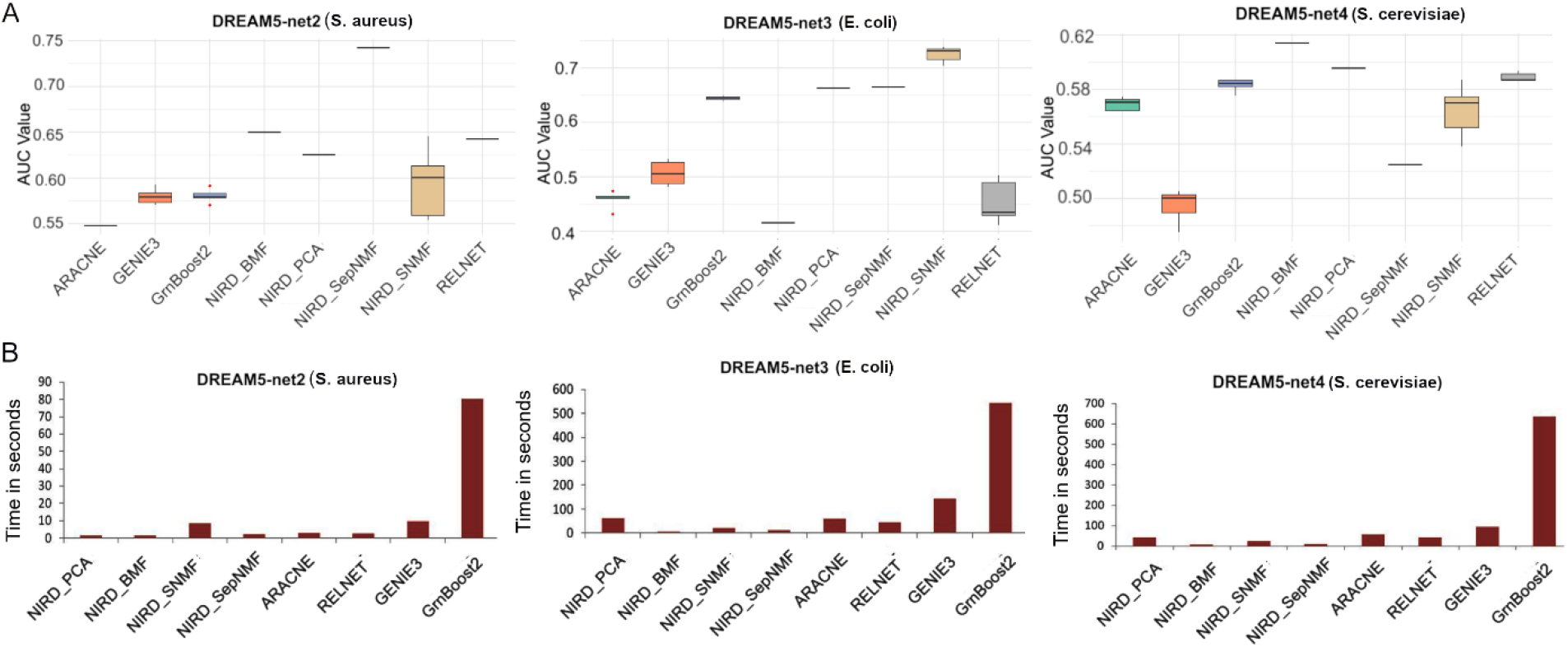
Evaluation of different network inference methods using bulk gene-expression profiles from DREAM 5 challenge. A) The values of Area under curve (AUC) shown as box-plot summarising the performance of different methods Dream 5, net2, net3 and net4 gene-expression datasets. Here performance of NIRD with four different types of matrix factorisation is also compared. For NIRD_BMF the matrix factorisation called binary factorisation method(BMF) was used. Similarly the names of NIRD_SepNMF, NIRD_PCA, NIRD_SNMF mention their corresponding matrix factorisation methods used. B)Time taken by different methods for mentioning expression datasets.

### NIRD enables better inference of gene-regulatory network using single-cell expression profiles

The pattern of single-cell transcriptomic data is drastically different from that of a bulk expression profile. Therefore, to test the robustness of the NIRD approach for single-cell expression, we benchmarked it against other methods using single-cell transcriptomic profiles of mouse embryonic stem cells [1,11]. In the evaluation using the literature curated gold-set of interaction [12] NIRD with PCA based dimension had best performance followed by NIRD with PMF based matrix factorisation (Fig. 3A). ARACNE and GRNBoost2 were the worst performers in terms of area under curve (AUC) for mESC single cell data (see Methods). We also used protein-protein interaction (PPI) information [13,14] to evaluate different network inference methods like Chen et al. [1]. NIRD based on PCA and SVD had 4-5 percent higher AUC level than GENIE3 (Fig. 3B). However, the available gold set interaction and PPI might have an incomplete set of gene-interactions. Therefore, we also used another measure based on the overlap among inferred networks from two scRNA-seq profiles of mESCs, with different batch effects and technical biases. If a method infers a gene network that is closer to true regulatory interactions, then, irrespective of technical bias, the predicted networks should have high overlap with each other. The network inferred using mESC single-cell transcriptomes generated using two different protocols (SMARTseq and Drop-seq) [11] were overlapped for evaluating different methods. NIRF, using PCA and PMF matrix factorisation, and ARACNE had substantially better performance than GENIE3 and GRNBoost2 (Fig. 3C). This is an important result, as it highlights that naive tree-based ensemble methods are unstable and highly sensitive to noise and bias in single-cell profiles. Whereas, an NIRD-based approach avoids such instability and noise by selecting only the top basis vectors after matrix factorisation, such as selecting only the top principal components in PCA-based dimension reduction. In addition, the NIRD-based approach required less computation time for single-cell transcriptomic profiles than tree-based ensemble methods (Fig. 3D).

**Figure 3.**
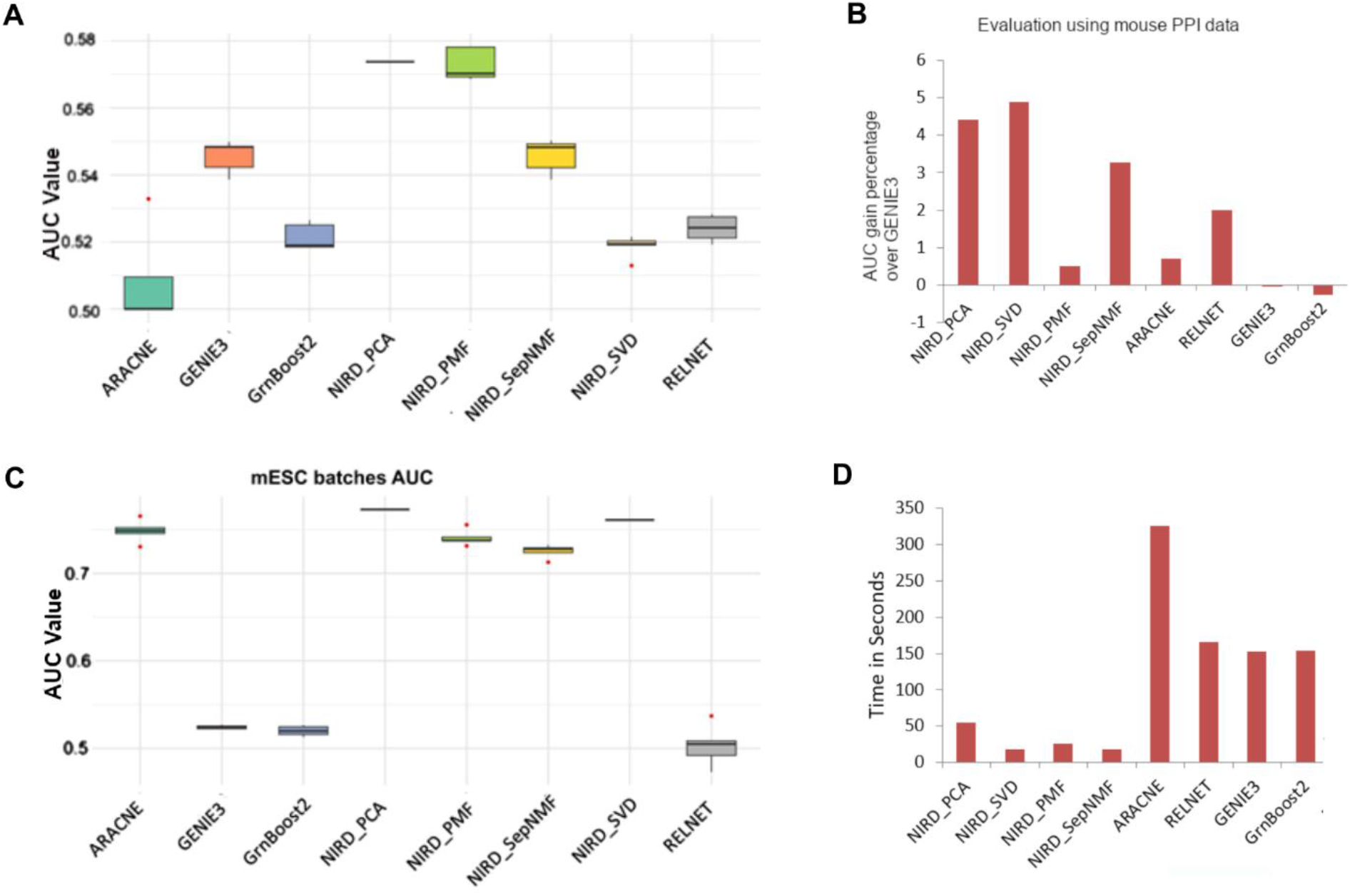
Evaluation of different network inference method using single-cell profiles of mouse embryonic stem cells. A) The box-plots show Area under curve (AUC) summarising the performance of different methods based on comparison with Gold set of gene regulatory interactions. B) The evaluation of inferred networks using mESC using known protein-protein interactions. C) AUC showing overlap of gene-interactions predicted using two datasets of mESC single-cell expression profiles using two different protocols. D) Time taken by different methods to infer gene-interaction network for mESC using the expression of 2000 genes.

### NIRD reveal cell-type specific regulatory differences between disease and normal cells

An application of gene-network inference using single-cell transcriptomes is to reveal influential genes from inferred networks across various cell types and individuals. For this purpose researchers can compare the network based importances of genes in inferred networks from diseased and normal samples. However, the first criterion for reliable results from such comparisons is that the inferred network for a cell type should remain largely consistent across samples from different individuals or across conditions and batch effects. At the same time, there should be relevant disease-specific changes in the importance of genes in inferred gene networks. We first evaluated network inference methods using the single-cell transcriptome of pancreatic beta cells of old and young individuals published by Enge et al [15], to check about their consistency in inferred interaction. GENIE3 and GRNBoost2 had substantially lower performance than the NIRD method, based on both the PPI set and AUC-based overlap of inferred networks for old and young pancreatic beta cells (supplementary Fig. S1).

Further we applied NIRD on single-cell profiles of cells from articular cartilage collected from normal individuals and patients with osteoarthritis (OA) and published by Fan et al (2024). We inferred and compared gene-networks for normal individuals and OA patients preHTC and HTC cells, as Fan et al. [16] reported preHTC and HTC cells to be potentially OA-critical cell types (Fig. 4). The performance of NIRD was superior to other tested network inference methods for single cell transcriptome profiles of HTC and preHTC cells, when evaluation was done using PPI dataset (Fig. 4A). Overall, NIRD shows superiority in providing consistency in inferred gene-network for a cell-type in comparison to other tested methods (supplementary Fig. S2).

**Figure 4.**
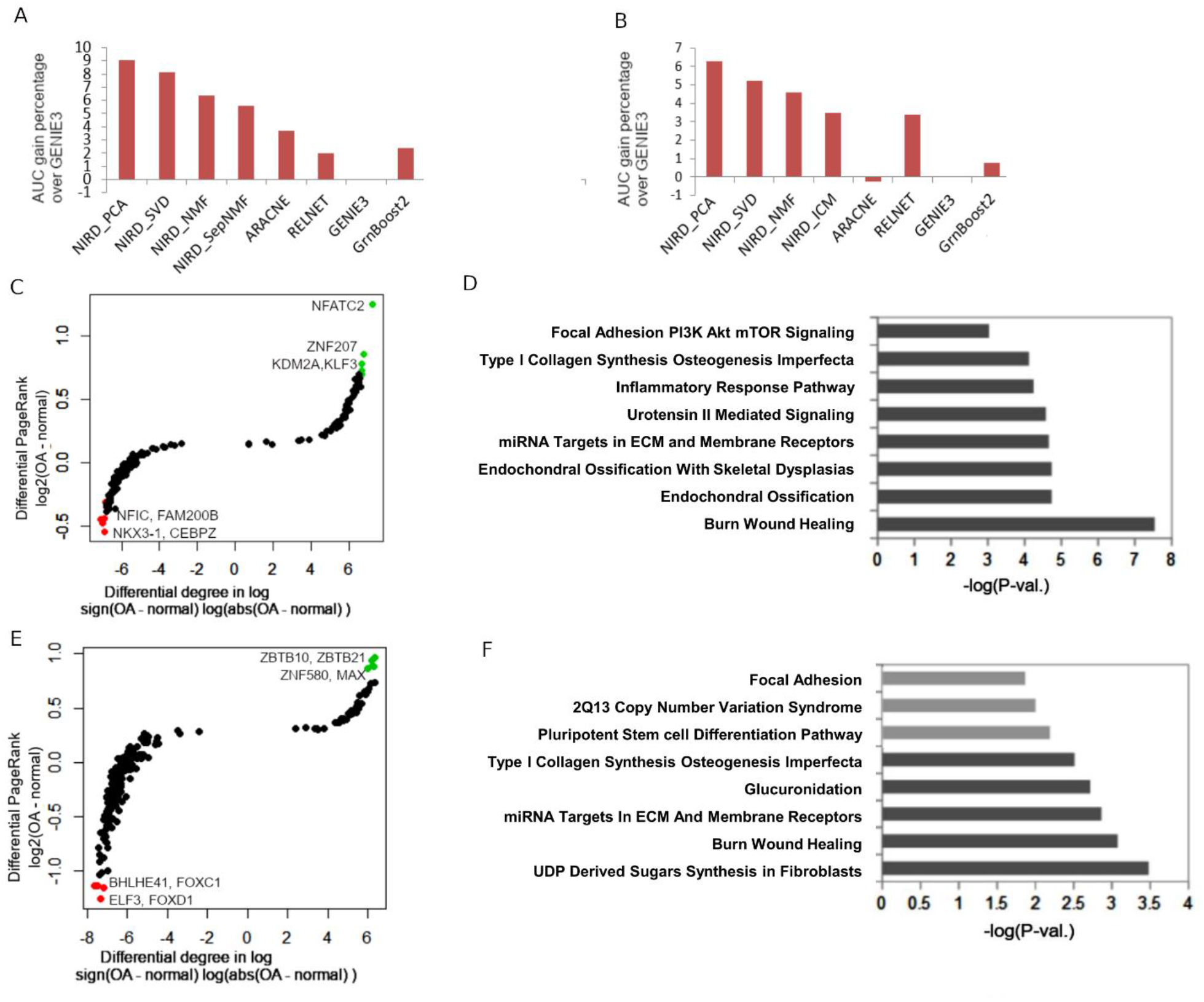
Analysis of inferred network using single-cell transcriptome profiles of cells from normal and arthritic patients. A) The evaluation of performance using known protein-protein interaction data. The AUC gains are shown here for different gene network inference methods applied to the single-cell transcriptome of the HTC cell. B) The performance in terms of AUC gain for different methods applied to single-cell transcriptome of preHTC cell C) The scatter-plot showing differential pageRank and differential degree of regulator nodes (transcription factors) in the inferred network using HTC cells from osteoarthritis patients (OA) and normal individuals. D) The enriched functions in top 30 genes with the highest positive differential PageRank in the inferred network for Osteoarthritis HTC compared to normal HTC. E) The plot shows differential pageRank and differential degree of transcription factors in the inferred network using preHTC single-cell transcriptome. F) The enriched functions in top 30 genes with the highest positive differential pageRank in comparison of inferred networks for Osteoarthritis preHTC versus normal preHTC.

Further comparison of gene degree and pageRank in inferred gene networks for normal and OA patients revealed top regulators that affect OA development. The transcription factors that showed a higher increase in pageRank in the inferred gene network for HTC single cells with OA compared to normal are NFATC2, ZNF207, KDM2A, and KLF3. While KDM2A has been previously linked to OA [17], our results suggest that HTC may be the cell type through which it exerts a greater effect on OA development. NFATC2 [18] and KLF3[19] have also been linked to inflammation and osteoarthritis previously. While the transcription factors with higher pageRank in normal sample HTC cells compared to OA include CEBPZ, NFIC, NKX3.1 and FAM200B (Fig. 4C). The top 8 functional terms enriched for top 50 genes with higher pageRank for network inferred from HTC cells in OA patients included ‘Burn Wound Healing’, ‘inflammatory response’ and Urotensin II mediated signalling, Focal adhesion PI3K_Akt_mTOR Signalling and endochondral ossification. Osteoarthritis is often called “a wound that does not heal” as tissue healing and repair processes get stalled in the inflammatory phase [20](Fig. 4D). Similarly, Urotensin II is known to be an inflammatory cytokine; however our results reveal that HTC cells are substantially affected by Urotensin II mediated pathways at OA joints.

For pre-HTC cells, the transcription factors with higher pageRank in the network inferred for the OA sample are ZBTB10, ZBTB21, ZNF580, and MAX (Fig. 4E). For the gene network in normal pre-HTC cells, the TFs with higher pageRank than in the OA network included ELF3, FOXD1, BHLHE41, and FOXC1. The top functional terms enriched for top 50 genes with higher pageRank for network inferred for preHTC of OA patients include ‘Burn Wound Healing’, ‘Glucuronidation’ and ‘UDP derived sugar synthesis in fibroblast’ (Fig. 4F). The Glucornidation [21,22] as such adds glucuronic acid to a substrate, its direct role in arthritis through preHTC cells have not been reported (Fig. 4F). It could be the effect of drugs being taken by the OA patient or there could be an unknown mechanism involved which Glucornidation affects arthritis. UDP-derived sugar synthesis leads to the production of glycosaminoglycans, which are essential components of the extracellular matrix [23]. Thus, for single-cell transcriptomes of synovial cells, NIRD demonstrated consistency across large-scale network inference, highlighted relevant genes, and enabled the identification of important pathways in a cell-type-specific manner.

### NIRD using RNA velocity

The framework of NIRD also allows using RNA-velocity and expression of genes simultaneously such that one can estimate the direct non-linear effect of regulators on poising and activation of genes (Methods and Material, Fig. 5A). However, quantification of dependency between expression of regulators and RNA-velocity of target genes using the approach of dimension reduction has rarely been reported before hence we performed an evaluation of NIRD. For brevity, we call the NIRD implementation using both RNA velocity and gene expression NIRD-expr+velo, and the simple NIRD implementation using only gene expression NIRD-expr-only. We applied NIRD-expr+velo and NIRD-expr-only on single-cell transcriptome profiles of human embryonic stem cells (hESCs)[24].

**Figure 5.**
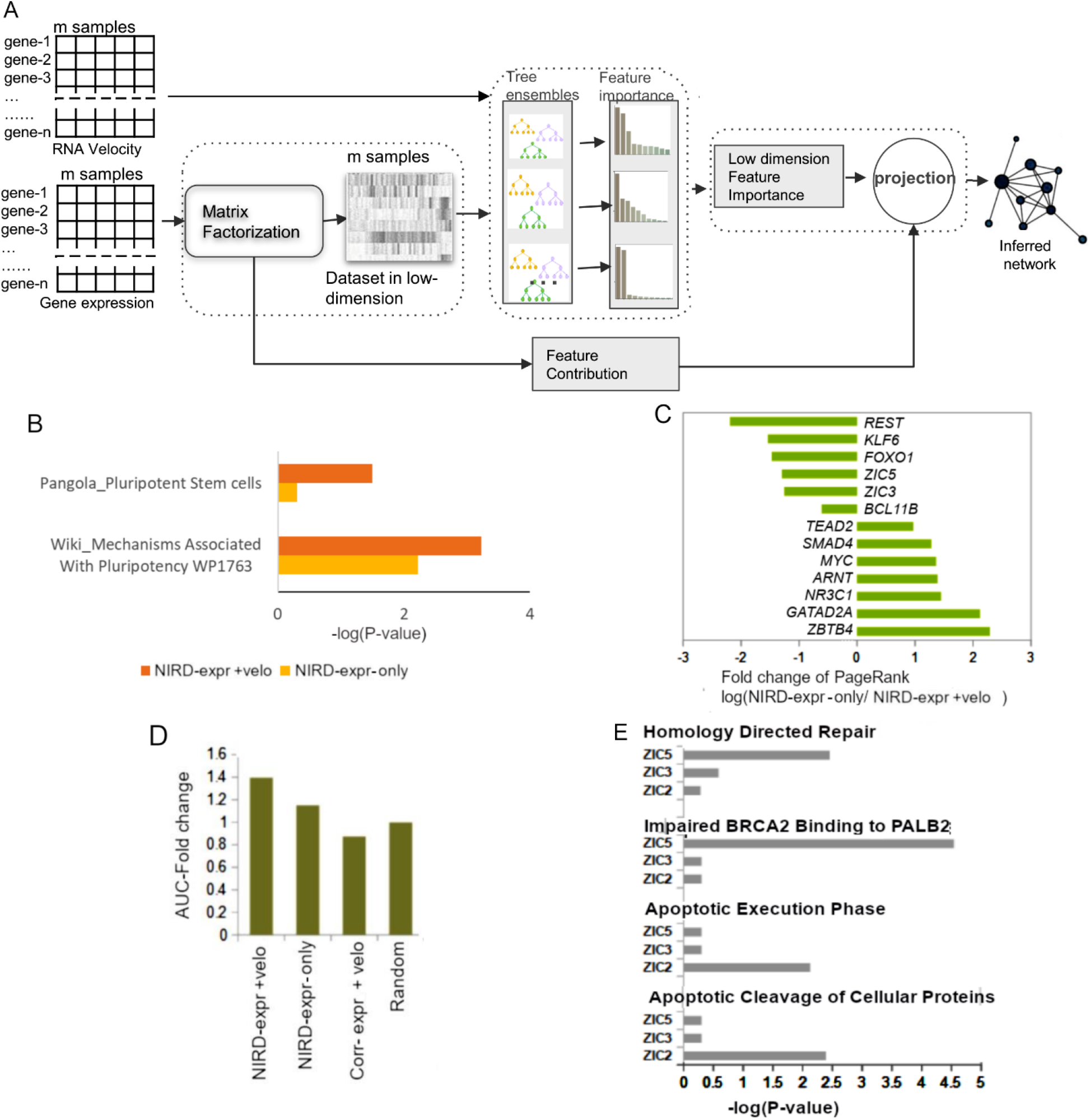
Evaluation of the approach of using RNA-velocity for gene-network inference in reduced dimension for primed human embryonic stem cells (hESC). A) Flowchart of the algorithm of using RNA-velocity and expression together with our approach of network inference in reduced dimension (NIRD) B) Comparison of enrichment for pathways of top genes with highest pageRank in gene-networks inferred using two approaches: NIRD-expr+velo which uses both RNA-velocity and expression of genes and NIRD_expr-only for inferring dependency network among genes using their expression only. C) The fold change in pageRank of regulating transcription factors in the gene-network inferred by NIRD-expr+velo and NIRD_expr-only. D) evaluation of performance of different network inferences for predicting genes being regulated by ZIC3 : NIRD-expr+velo, NIRD-expr-only, and network inference using correlation between velocity and expression (corr-expr+velo). The positive gold set consisted of genes with a ZIC3 ChIP-seq peak at their promoters and a 3-fold increase in expression upon ZIC3 knockdown in primed hESCs. E) Functional terms enriched for NIRD-velocity based predicted genes for being regulated by ZIC3, ZIC2 and ZIC5 in primed hESC.

For hESC, we compared the enrichment of ontology terms of the top 50 genes with the highest pageRank in networks inferred by NIRD-expr+velo and NIRD-expr-only. We found more significant enrichment of pluripotency terms for top 50 genes from NIRD-expr+velo than NIRD-expr-only (Fig. 5B). Further, we found TFs known to have an important role in pluripotency and priming for early differentiation of hESCs such as ZIC3, ZIC5, FOXO1, KLF6, REST had higher pageRank in the network inferred by NIRD-expr+velo (Fig. 5C). Where only a few TFs known for pluripotency like SMAD4, TEAD2 and Myc had higher pageRank in the gene-network inferred using NIRD-expr-only.

Further, we choose a transcription factor, ZIC3, which is known to play an important role in primed hESC cells. In primed human embryonic stem cells (hESC) we chose those genes as directly regulated by ZIC3 which had ZIC3 ChIP-seq peaks (Hossain et al. 2024)and showed substantial change(fold change > 3) in expression upon its knock down (Fig. 5D). We found that coverage of directly regulated genes by ZIC3 as quantified by AUC was better in a network estimated by NIRD-expr+velo than by NIRD-expr-only and random null model. We even inferred another network by finding correlation of every gene RNA-velocity with expression of other genes (named as corr-expr+velo in Fig. 5D). However, we found that correlation based network inference were not able to highlight directly regulated genes of ZIC3 over other genes, as its performance in terms of AUC was comparable to random null model (with random connections among genes). Whereas AUC for NIRD-expr+velo was 1.4 times better than random null model. Such a result suggests that combining NIRD with RNA velocity and expression can reveal directly influenced target genes and thus help elucidate the function of a transcription factor. Therefore we further investigated and compared the roles of TFs ZIC3, ZIC5 and ZIC2 predicted using NIRD-expr+velo. We choose genes in the top five percentile score of interaction and having correlation of their RNA-velocity above 0.1 with expression of ZIC3, ZIC2 and ZIC5. We found differences in enrichment of ontology terms for top interacting genes for ZIC2, ZIC3 and ZIC5 in a network inferred by NIRD-expr+velo (Fig. 5E). Apoptosis related functions were more enriched for top genes with the highest interaction score for ZIC2 in comparison to top targets of ZIC3 and ZIC5 (Fig. 5E). For target genes of ZIC5 the top enriched function terms were related to homology directed repair function especially involving BRCA1 and BRCA2. ZIC2 is known to be involved in proliferation and reducing apoptosis in ESC as acute ablation of ZIC2 causes death in differentiating ESC[25]. However we could not find published literature related to association between ZIC5 regulation of genes involved in homology directed repair. Nevertheless, overall, NIRD used with RNA-velocity could capture the regulatory dependencies among genes in primed hESC as well as facilitate prediction of function of TFs.

## Discussion

Low-rank approximation of data matrices, along with Machine Learning, is a topic of extensive research. Low-rank approximation provides an efficient, compact representation of multidimensional vectors by embedding high-dimensional data, rescuing analytics by mitigating noise, unfolding latent relations, and handling sparsity. Our results clearly show that, for large, sparse single-cell transcriptomic datasets, we may not achieve reliable consistency with methods like GENIE3 and GRNBoost2, which rely solely on machine learning and do not handle the noise and complexity of high-dimensional data. The NIRD framework allows low-rank approximation using multiple dimension-reduction methods, as long as we can estimate the contribution of the original features to the lower-dimensional basis vectors. However, for each combination of dimension reduction method and ensemble-based tree method, it may be necessary to benchmark across different types of datasets. The current NIRD framework using randomForest or extraTree can also be applied to other types of single-cell profile datasets [26]. However, in this study, we have focused only on analysing single-cell transcriptomic and RNA velocity profiles using NIRD.

A method proposed by Li et al. [27] performs low-dimensional embeddings over genes and cells for network inference. However their method further uses two multi-layer perceptrons (MLPs) over gene and cell embedding to find gene-TF regulatory interactions [27]. Previously, a few other methods used projection of gene expression into lower dimensions for faster inference of large gene networks; however, most of them rely on the notion of “guilt by association” [28] [29]. They assume that genes that are co-localised in lower dimensions are more likely to regulate each other. However, the major issue with methods based on “guilt by association” is that they highlight redundant co-regulation based on association and often ignore the effects of regulators and their combinatorics. It has already been shown that changes in gene expression in different disease conditions highlight only the effect, not the causality [30]. In such cases the ‘guilt by association’ approach could wrongly prioritise indirect relationships between affected genes with similar response or co-expression and ignore the causal genes . On the other hand, GENIE3 and NIRD-like approaches rely on non-linear effects of regulator genes, which mimic the natural combinatorial causal effects of transcription factors on gene activity.

Our analysis using gold data and mESC datasets shows that data sparsity contributes to instability in tree-based ensemble methods. The NIRD-based approaches address these bottlenecks by selecting the top variation-driving vectors during matrix factorisation. We find that NIRD consistently captures a larger network, including several known and new targets, as well as pathways, in a cell-type-specific manner. Analysis of single-cell transcriptomes from HTC and preHTC using NIRD suggests a potential role for a few TFs previously unknown to be linked to OA. Our results indicate that ZNF207 is likely to be important for OA-related inflammation through its activity in HTC cells. Similarly, ZBTB10, ZBTB21, ZNF580, MAX contribute to OA associated inflammation in preHTC cells. Previously, Zhang et al. reported the involvement of ZBTB10 in rheumatoid arthritis and its regulation by miR-361-5p expressed in synovial tissues. However, for osteoarthritis the role of ZBTB10 still needs to be thoroughly studied. The concentration of MAX (heterodimer partner of MYC) is mainly in the nucleus of proliferative chondrocytes and decreases as chondrocytes mature[31]. Thus MAX[31] could have a direct effect on the reduction of differentiation and proliferation of pre-chondrocytic prehypertrophic (preHTC) cells, which are actively involved in OA.

In most available single-cell transcriptomic profiles, time-series datasets are not available; hence, inferring causality in gene regulation is not trivial in such cases. Prior knowledge of trajectory directionality is useful for minimising spurious predictions. Our data show that using NIRD with RNA velocity can resolve the problem of inferring causality to some extent, as it emphasises direct target genes. We observe that NIRD-expr+velo was superior to NIRD-expr_only in detecting directly regulated genes by ZIC3, as highlighted by ChIP-seq peaks and knockdown-based experimental observations. Such results are quite encouraging for researchers keen to predict future response of cells. A few studies have used GRNBoost2 with additional knowledge of possible targets of TFs, and we believe this prior knowledge can be useful in pruning and selecting biologically relevant targets(Bravo González-Blas et al. 2023). In the current study, we have focussed on NIRD based core network-inference and robust evaluation using statistical criteria. We expect that both prior knowledge, additional target information and features can also be added to NIRD to make it a more biologically accurate method for down-stream analysis.

## Conclusion

The study here proposes an efficient approach of finding non-linear dependencies between large number features using their scores across multiple observations. Thus the proposed approach is ideal for inferring large regulatory networks, especially gene-interaction networks, which was demonstrated here using bulk and single-cell transcriptome profiles. The proposed method outperformed multiple methods on the benchmark using a gold-standard of known interactions, protein-protein interaction data, and consistency despite batch effects. The consistency of network inference across batch effects achieved by NIRD was demonstrated across 3 single-cell transcriptomic datasets. The consistency of the inferred network produced by the proposed approach helped highlight relevant regulators that may be involved in osteoarthritis development, given the diverse cell types present in articular cartilage. A further strength of the proposed approach is its flexibility in combining RNA velocity with gene expression to improve the prediction of direct gene regulators. The proposed method, using RNA velocity on single-cell profiles of human embryonic stem cells, was superior at identifying direct target genes of ZIC3. The proposed method, combined with RNA velocity, also revealed direct targets of ZIC2 and ZIC5 and their effects on different functions.

## Material and Methods

### Inferring non-linear dependencies after dimension reduction

Most network inference algorithms do not work efficiently in high-dimensional spaces due to the size of the data and the curse of dimensionality. To find non-linear dependencies among features using their scores at data points, we have relied on tree-based ensemble methods (Random Forest and Extra Trees). However, unlike Genie3[4] we do not try to model the expression of a gene using expression of other genes. We first calculate a lower-dimensional representation of cell expression profiles, mapping each data point or cell to a set of basis vectors. We try to model expression of every gene using the basis vectors. Further, we calculate the loading of each gene onto the basis vector. Hence, given the expression matrix *X* (dimension *m* × *n*) with *m* cells (data-point) and *n* genes (features), we factorise it to matrices A (dimension *m* × *k*) and Y (dimension *k* × *n*) such that columns of matrix *A* contains the basis vector. Hence it can be written as

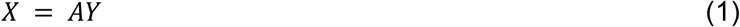

We model expression of genes using the elements of basis vectors in *A* using a tree ensemble-based machine learning method. Such that for gene *j*, the corresponding column in the expression matrix *X* is modelled using columns in matrix *A*. It can be written that jth column (dimension *m* × 1) of matrix *X*

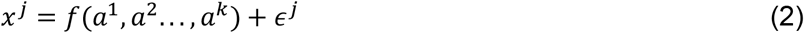

Here *a*^*i*^ represents the *i*th column vector (dimension *m* × 1) of matrix *A* and *ϵ*^*j*^ represents a vector of noise corresponding of expression of gene *j*. While *f*() represents the modelling function, which in our case is a tree-based non-linear regression model. When we use tree-based methods (e.g., Random Forests or Extra Trees), we can estimate feature importance. Thus, here we estimate feature importance of every basis vector *i* as *F*_*i*_ ^*j*^ while modelling the expression of gene *j* across m samples (cells). We project the feature importances back to the original features (genes) using a linear approach. Hence, we calculate the feature importance of the original feature *l* (gene expression *x*^*l*^) while modeling the expression of gene *j* as *w*_*lj*_,using the equation

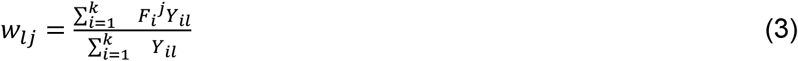

Where *Y*_*il*_ is the element in the coefficient matrix *Y* corresponding to *ith* row and *lth* column. Using the method described here we calculate all original features (genes or regulator) importance while modeling expression of each gene. The original feature importance *w*_*lj*_ is considered here as weight of edge between genes *l* and *j*.

Here for demonstration purpose we used 14 matrix factorisation approaches which include : Principal component analysis (PCA), kernel PCA, singular value decomposition (SVD), non-negative matrix factorisation (NMF), Bayesian decomposition(BD), Binary matrix factorisation (BMF), Iterated Conditional Modes (ICM) NMF, Fisher Nonnegative Matrix Factorization for learning Local features (LFNMF), sparse non-negative matrix factorisation (SNMF), probabilistic matrix factorisation (PMF), Penalised matrix factorisation for constrained clustering (PMFCC), separable NMF (SepNMF). ENMF and KLD-NMF. Each matrix factorisation method has its pros and cons. While principal components analysis and vector quantisation can be understood as matrix factorisation techniques for low-rank matrix approximation/estimation under different constraints. Different types of constraints used by matrix factorisation methods can have distinct effects on representational properties.

Principal components analysis imposes only an orthogonality constraint, resulting in a highly non-redundant representation that uses cancellations to generate variability [32]. Nonnegativity is a useful constraint for matrix factorisation that can learn a partial representation of the data [33]. The nonnegative basis vectors learned by methods like NMF, SepNMF, ENMF, and KLD-NMF utilise distributed and sparse combinations to achieve expressiveness.

### Using NIRD with RNA velocity

NIRD can also be used by providing it with a matrix of RNA-velocities and a matrix with gene-expression values. Given the RNA velocity matrix *R* where each row represents a gene and every column is for a single-cell, we try to model the elements in a row using the basis vectors of A. As described above, here A is the reduced dimension representation of the gene-expression matrix. Hence if gene expression matrix of all genes or selected genes such as transcription factor is provided as matrix *X*, we factorise it using matrix factorisation as shown in equation (1). The column vectors of matrix *A* is used to model RNA-velocity of genes. Such that for gene *j* the vector RNA velocity or the jth row in matrix *R* (*r*^*j*^) can be written as

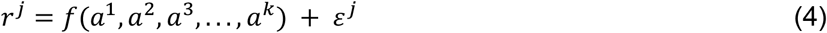

Where *a*^*i*^ represents the *i*th column vector (dimension *m* × 1) of matrix *A* and *ε*^*j*^ represents vector of noise corresponding RNA velocity of gene *j*. As mentioned above we estimate the feature importance of actual features (gene expression) while modelling RNA velocity of gene *j* . Similarly we also calculate feature-importances for all genes which have their RNA velocity in matrix *R*. The feature importances here correspond to the weights of the dependencies between two genes; in other words, they estimate the effect of gene expression on the RNA-velocity of another gene. We estimated RNA-velocity using single-cell RNA-seq according to the pipeline described by Velocyto [34] tool.

### Evaluation of different methods

We used the approach of Area under curve (AUC) to evaluate different methods. For this purpose, we calculated the area under the Receiver operating characteristic (ROC) curve by comparing edges in the inferred network with those in the ground truth. For the Receiver operator curve, the X-axis showed the number of edges sorted by their weights instead of the false-positive rate. The reason for this approach is that, in the absence of exhaustive true-positive edges, it is not trivial to obtain false-positive interactions. We calculated AUC under ROC using all the possible edges provided by the network inference method. For Dream 5 challenge dataset ground truth (gold-set) of interactions was available. For networks inferred using mESC single-cells we used the benchmark set of known interactions published in literature and curated by Meisig and Bluthgen [12]. We also used an evaluation method in which we inferred the network using two batches of single-cell transcriptomes from the same cell type. It is based on the notion that if the network inferred represents true interactions, then we would predict similar edge weights, irrespective of batch effects in single-cell profiles for the same cell type. To evaluate the consistency of the inferred networks for mESC we used single-cell profiles from using 2 different protocols (SMARTseq and Drop-seq) by Ziegenhan et al. [11]. We also used known protein-protein interaction data to evaluate the network inference methods.

### Evaluation based on transcription factor ChiP-seq and knock-down in hESC

In order to check whether NIRD using RNA-velocity and gene-expression provided better inference of direct dependencies among genes, we used a TF ZIC3 chip-seq profile and gene-expression data from its knock down in primed hESC(Hossain et al. 2024). We aligned ZIC3 ChIP-seq reads to human genome (version hg19), the peaks were called using DFilter[35]. The significant peaks (DFilter p-value < 1E-8) of ZIC3 ChIP-seq were then intersected with the RefSeq gene promoter regions within 750 bp of transcription start sites (hg19 version). The genes with ZIC3 peaks in their promoters and substantial changes in expression upon ZIC3 (fold change > 3, up or down) were chosen as positive (gold set). In order to estimate performance of coverage of positives (genes regulated by ZIC3) using inferred network, we used only the interaction strength of genes with ZIC3 in four different gene-networks inferred using NIRD-expr+velo, NIRD-expr-only, random and correlation between expression and RNA-velocity. For each method, the interaction strength with ZIC3 was used to calculate the ROC for coverage of the gold-set interactions with ZIC3. Finally the AUC gain for different methods was calculated with respect to AUC of the random network (shown in Fig. 5D).

## Data sources

The single-cell expression profile of mESC from different protocols was generated by Ziegenhan et al. and is available in the GEO database (GEO id: GSE75790). The benchmark for mESC network inference curated by Mei sig and Bluthgen was downloaded from github ([GitHub-johannesmg/benchmark_gene_regulation_mesc](https://github.com/johannesmg/benchmark_gene_regulation_mesc)). Single-cell transcriptome of cartilage tissue from OA and normal individuals were downloaded from GEO dataset (GSE255460). The single-cell expression profiles of pancreas of old and young individuals were published by Enge et al [15] and are available at GEO dataset(GEO ID:GSE81547). The single-cell transcriptomes of hESC were adapted from the GEO dataset (GSE75748). The raw sequencing files for ChIP-seq of ZIC3 and input control in hESC were downloaded from the SRA dataset corresponding to the GEO ID : GSE270784(Hossain et al. 2024).

## Availability of code

The code for NIRD can be downloaded from the Github link https://github.com/reggenlab/NIRD The Github repository would be made public after acceptance of manuscript The codes and results can be downloaded from http://reggen.iiitd.edu.in:1207/NIRD/ or http://reggen.iiitd.edu.in:1207/NIRD/codes/

For access User id: reggen and password unipath@123

### Abbreviations

NIRD: Network inference in reduced dimension
OA: Osteoarthritis
PCA: Principal component analysis
SVD: singular value decomposition
NMF: non-negative matrix factorisation
BD: Bayesian decomposition
BMF: Binary matrix factorisation (BMF)
ICM: Iterated Conditional
SNMF: sparse non-negative matrix factorisation
PMF: probabilistic matrix factorisation
PMFCC: Penalised matrix factorisation for constrained clustering
SepNMF: separable non-negative matrix factorisation
KLD: 
AUC: area under curve
TF: transcription factors
PPI: protein-protein interaction
ChIP: chromatin immunoprecipitation
HTC: hypertrophic chondrocytes
preHTC: prehypertrophic chondrocytes

## Author Contributions

Conceptualisation : V.K.1, V.K.2, Method Implementation: I. P. J, A. G. M., Data curation and Analysis: V.K.2, I.P.J., A.G.M. Supervision : V.K.1 Review and writing: V. K.1 and K. N.N.

## References

1. Chen S, Mar JC. Evaluating methods of inferring gene regulatory networks highlights their lack of performance for single cell gene expression data. BMC Bioinformatics. BioMed Central; 19:1–212018;

2. Pratapa A, Jalihal AP, Law JN, Bharadwaj A, Murali TM. Benchmarking algorithms for gene regulatory network inference from single-cell transcriptomic data. Nature Methods. Nature Publishing Group; 17:147–542020;

3. Marbach D, Costello JC, Küffner R, Vega N, Prill RJ, Camacho DM, et al. Wisdom of crowds for robust gene network inference. Nature methods. 9:7962012;

4. Huynh-Thu VA, Irrthum A, Wehenkel L, Geurts P. Inferring Regulatory Networks from Expression Data Using Tree-Based Methods. PLoS ONE. 5:e127762010;

5. Moerman T, Aibar SS, Bravo G-BC, Simm J, Moreau Y, Aerts J, et al. GRNBoost2 and Arboreto: efficient and scalable inference of gene regulatory networks. Bioinformatics (Oxford, England). Bioinformatics; 2019; doi: 10.1093/bioinformatics/bty916.

6. Aibar S, González-Blas CB, Moerman T, Huynh-Thu VA, Imrichova H, Hulselmans G, et al. SCENIC: single-cell regulatory network inference and clustering. Nature Methods. Nature Publishing Group; 14:1083–62017;

7. Dong J, Li J, Wang F. Deep Learning in Gene Regulatory Network Inference: A Survey. IEEE/ACM transactions on computational biology and bioinformatics. IEEE/ACM Trans Comput Biol Bioinform; 2024; doi: 10.1109/TCBB.2024.3442536.

8. Chevalley M, Roohani YH, Mehrjou A, Leskovec J, Schwab P. A large-scale benchmark for network inference from single-cell perturbation data. Communications Biology. Nature Publishing Group; 8:1–182025;

9. Margolin AA, Nemenman I, Basso K, Wiggins C, Stolovitzky G, Favera RD, et al. ARACNE: An Algorithm for the Reconstruction of Gene Regulatory Networks in a Mammalian Cellular Context. BMC Bioinformatics. BioMed Central; 7:1–152006;

10. Faith JJ, Hayete B, Thaden JT, Mogno I, Wierzbowski J, Cottarel G, et al. Large-Scale Mapping and Validation of Escherichia coli Transcriptional Regulation from a Compendium of Expression Profiles. PLOS Biology. Public Library of Science; 5:e82007;

11. Ziegenhain C, Vieth B, Parekh S, Reinius B, Guillaumet-Adkins A, Smets M, et al. Comparative Analysis of Single-Cell RNA Sequencing Methods. Molecular cell. Mol Cell; 2017; doi: 10.1016/j.molcel.2017.01.023.

12. Meisig J, Blüthgen N. The gene regulatory network of mESC differentiation: a benchmark for reverse engineering methods. Philosophical transactions of the Royal Society of London Series B, Biological sciences. Philos Trans R Soc Lond B Biol Sci; 2018; doi: 10.1098/rstb.2017.0222.

13. Alanis-Lobato G, Andrade-Navarro MA, Schaefer MH. HIPPIE v2.0: enhancing meaningfulness and reliability of protein–protein interaction networks. Nucleic Acids Research. 45:D4082016;

14. Szklarczyk D, Kirsch R, Koutrouli M, Nastou K, Mehryary F, Hachilif R, et al. The STRING database in 2023: protein–protein association networks and functional enrichment analyses for any sequenced genome of interest. Nucleic acids research. Nucleic Acids Res; 2023; doi: 10.1093/nar/gkac1000.

15. Enge M, Arda HE, Mignardi M, Beausang J, Bottino R, Kim SK, et al. Single-Cell Analysis of Human Pancreas Reveals Transcriptional Signatures of Aging and Somatic Mutation Patterns. Cell. Cell; 2017; doi: 10.1016/j.cell.2017.09.004.

16. Yue Fan, Xuzhao Bian, Xiaogao Meng, Lei Li, Laiyi Fu, Yanan Zhang, Long Wang, Yan Zhang, Dalong Gao, Xiong Guo, Mikko Juhani Lammi, Guangdun Peng, Shiquan Sun. Unveiling inflammatory and prehypertrophic cell populations as key contributors to knee cartilage degeneration in osteoarthritis using multi-omics data integration. Annals of the Rheumatic Diseases. Elsevier; 83:926–442024;

17. Fan Y, Bian X, Meng X, Li L, Fu L, Zhang Y, et al. Synovial Mesenchymal Stem Cell-Derived EV-Packaged miR-31 Downregulates Histone Demethylase KDM2A to Prevent Knee Osteoarthritis. Molecular Therapy - Nucleic Acids. Cell Press; 22:1078–912020;

18. Lv R, Du L, Bai L. RNF125, transcriptionally regulated by NFATC2, alleviates osteoarthritis via inhibiting the Wnt/β-catenin signaling pathway through degrading TRIM14. International immunopharmacology. Int Immunopharmacol; 2023; doi: 10.1016/j.intimp.2023.111191.

19. You M, Ai Z, Zeng J, Fu Y, Zhang L, Wu X. Bone mesenchymal stem cells (BMSCs)-derived exosomal microRNA-21-5p regulates Kruppel-like factor 3 (KLF3) to promote osteoblast proliferation in vitro. Bioengineered. Bioengineered; 2022; doi: 10.1080/21655979.2022.2067286.

20. Huston P. Why osteoarthritis of the knee is called “a wound that does not heal” and why Tai Chi is an effective treatment. Frontiers in Medicine. 10:12083262023;

21. {Ortutay Z, Polgár A, Gömör B, Géher P, Lakatos T, Glant TT, et al. Synovial fluid exoglycosidases are predictors of rheumatoid arthritis and are effective in cartilage glycosaminoglycan depletion. Arthritis and rheumatism. Arthritis Rheum; 2003; doi: 10.1002/art.11093.

22. Meunier CJ, Verbeeck RK. Glucuronidation of R- and S-Ketoprofen, Acetaminophen, and Diflunisal by Liver Microsomes of Adjuvant-Induced Arthritic Rats. Drug Metabolism and Disposition. Elsevier; 27:26–311999;

23. Zimmer BM, Barycki JJ, Simpson MA. Integration of Sugar Metabolism and Proteoglycan Synthesis by UDP-glucose Dehydrogenase. Journal of Histochemistry and Cytochemistry. 69:132020;

24. Chu LF, Leng N, Zhang J, Hou Z, Mamott D, Vereide DT, et al. Single-cell RNA-seq reveals novel regulators of human embryonic stem cell differentiation to definitive endoderm. Genome biology. Genome Biol; 2016; doi: 10.1186/s13059-016-1033-x.

25. Luo Z, Gao X, Lin C, Smith E, Marshall S, Swanson SK, et al. Zic2 is an enhancer-binding factor required for embryonic stem cell specification. Molecular cell. 57:6852015;

26. Baysoy A, Bai Z, Satija R, Fan R. The technological landscape and applications of single-cell multi-omics. Nature Reviews Molecular Cell Biology. Nature Publishing Group; 24:695–7132023;

27. Li S, Liu Y, Shen L-C, Yan H, Song J, Yu D-J. GMFGRN: a matrix factorization and graph neural network approach for gene regulatory network inference. Briefings in Bioinformatics. 25:bbad5292024;

28. Pindah W, Seman A, Nordin S, Said MSM. Review of dimensionality reduction techniques using clustering algorithm in reconstruction of gene regulatory networks. IEEE;

29. Kc K, Li R, Cui F, Yu Q, Haake AR. GNE: a deep learning framework for gene network inference by aggregating biological information. BMC Systems Biology. BioMed Central; 13:1– 142019;

30. Porcu E, Sadler MC, Lepik K, Auwerx C, Wood AR, Weihs A, et al. Differentially expressed genes reflect disease-induced rather than disease-causing changes in the transcriptome. Nature Communications. Nature Publishing Group; 12:1–92021;

31. Wang Y, Toury R, Hauchecorne M, Balmain N. Expression and subcellular localization of the Myc superfamily proteins: c-Myc, Max, Mad1 and Mxi1 in the epiphyseal plate cartilage chondrocytes of growing rats. Cellular and molecular biology (Noisy-le-Grand, France). Cell Mol Biol (Noisy-le-grand); 431997;

32. Vidal R, Ma Y, Sastry S. Generalized Principal Component Analysis. Springer;

33. Guo Y-T, Qin-Qin L, Chun-Sheng L. The rise of nonnegative matrix factorization: Algorithms and applications. Information Systems. Pergamon; 123:1023792024;

34. La Manno G, Soldatov R, Zeisel A, Braun E, Hochgerner H, Petukhov V, et al. RNA velocity of single cells. Nature. Nature Publishing Group; 560:494–82018;

35. Kumar V, Muratani M, Rayan NA, Kraus P, Lufkin T, Ng HH, et al. Uniform, optimal signal processing of mapped deep-sequencing data. Nature biotechnology. Nat Biotechnol; 2013; doi: 10.1038/nbt.2596.

