## supplementary Figures for "Inferring large networks with matrix factorisation to capture non-linear dependencies among genes using sparse single-cell profiles"

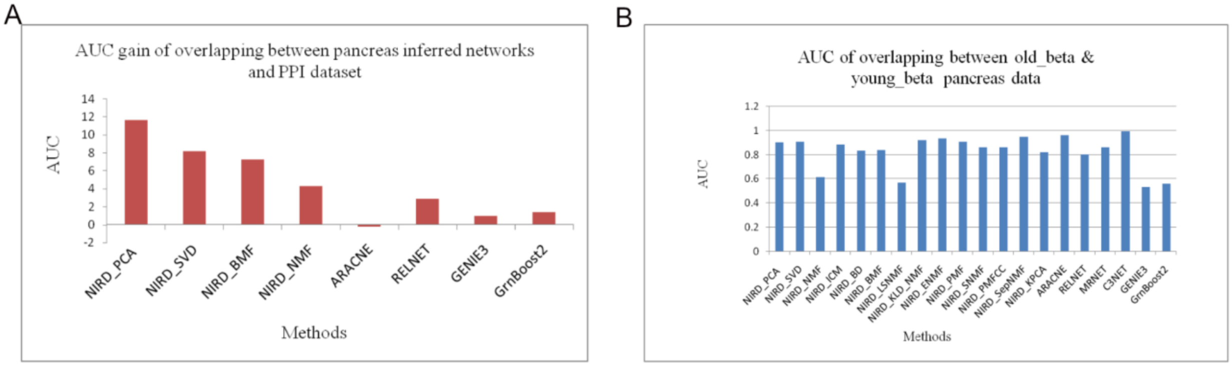


Figure S1 : The evaluation for network inference using single-cell transcriptome profiles of pancreatic beta cells from old and young.  A) performance was evaluated using protein-protein interaction data-set  B) performance was evaluated based on overlap between inferred networks from pancreatic beta cells from old and young individuals (Enge et al. 2017).

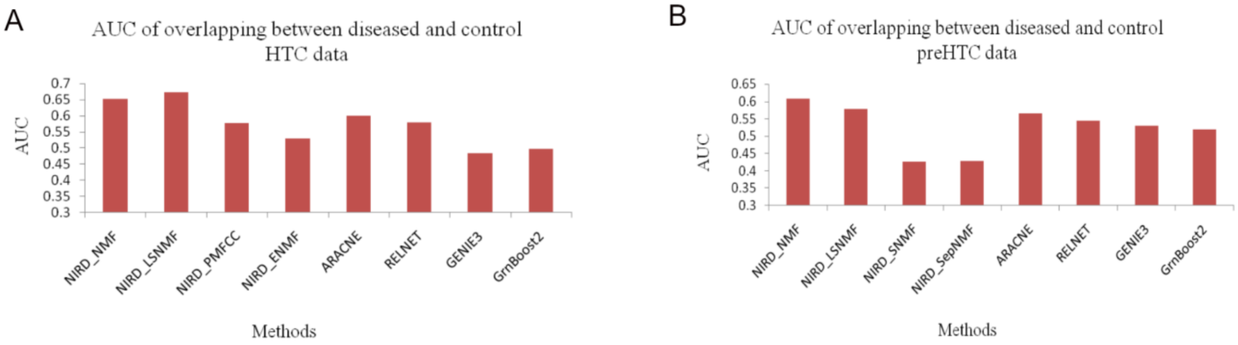


Figure S2:  Evaluation of performance for network inference using single-cell transcriptome profile of (A) HTC (hypertrophic chondrocytes) and (B) preHTC (prehypertrophic chondrocytes) cells for normal individuals and osteoarthritic patients. The evaluation was performed using the overlap between predicted networks from normal and OA cells.
